# A universal scaling method for biodiversity-ecosystem functioning relationships

**DOI:** 10.1101/662783

**Authors:** K.E. Barry, G.A. Pinter, J.W. Strini, K. Yang, I.G. Lauko, S.A. Schnitzer, A.T. Clark, J. Cowles, A.S. Mori, L. Williams, P.B. Reich, A.J. Wright

## Abstract

Global biodiversity is declining at rates faster than at any other point in human history. Experimental manipulations of biodiversity at small spatial scales have demonstrated that communities with fewer species consistently produce less biomass than higher diversity communities. However, understanding how the global extinction crisis is likely to impact global ecosystem functioning will require applying these local and largely experimental findings to natural systems at substantially larger spatial and temporal scales. Here we propose that we can use two simple macroecological patterns – the species area curve and the biomass-area curve – to upscale the species richness-biomass relationship. We demonstrate that at local spatial scales, each additional species will contribute more to biomass production with increasing area sampled because the species-area curve saturates and the biomass-area curve increases monotonically. We use species-area and biomass-area curves from a Minnesota grassland and a Panamanian tropical dry forest to examine the species richness – biomass relationship at three and ten sampling extents, respectively. In both datasets, the observed relationship between biodiversity and biomass production at every sampling extent was predicted from simple species-area and biomass-area relationships. These findings suggest that macroecological patterns like the species-area curve underpin the scaling of biodiversity-ecosystem functioning research and can be used to predict these relationships at the global scales where they are relevant for species loss.

---

Worldwide, drastic environmental changes are leading to biodiversity loss at regional and global scales^1–3^. Many predict that the rate of species loss will accelerate in the coming decades^4, 5^. Problematically, biodiversity supports many crucial ecosystem functions and services such as biomass production, carbon sequestration, and nitrogen cycling and retention^6–8^. That is, in small scale experiments, ecosystem functioning typically increases with increasing species richness^9–16^. In natural ecosystems, species richness appears to have an even larger effect on some functions like biomass production^17, 18^. Thus, ongoing species loss will likely have serious consequences for how ecosystems function^19, 20^. Yet the results from local-scale experiments and observational studies may have little relevance at the regional and global scales where species loss is occurring^21^. Understanding how to link these local-scale results to the regional and global scales where species loss is occurring is crucial for predicting the consequences of the global extinction crisis.

Here, we consider spatial extents ranging from 16 m^2^ to 400 m^2^ and predict that species richness may contribute more to biomass production with increasing sampling extent. This increase in the contribution of species richness to biomass production occurs because scaling the species richness-biomass relationship requires combining the species area curve and the biomass area curve (Figure 1). Species richness saturates with increasing area^22–30^ (Figure 1a). By contrast, biomass production increases monotonically with increasing area (hereafter, sampling extent) ^31^ (Figure 1a, Figure S1). Combining these two different scale-dependencies - a linear increase in biomass production and saturating species richness - results in an increase in the contribution of each additional species to biomass production with increasing sampling extent. Consequently, the slope of the species richness-biomass relationship will increase (Figure 1c). Alternatively, biomass production in biodiversity-ecosystem functioning studies is commonly standardized per unit area (e.g. biomass per m^2^, e.g., ^8, 11, 12, 14, 32–34)^. In per unit area terms, biomass production should be invariant to area (Figure 1d). This standardization means that each species represents a smaller contribution to biomass production (per unit area) as we increase the sampling extent (Figure 1e,f).

**Figure 1.**
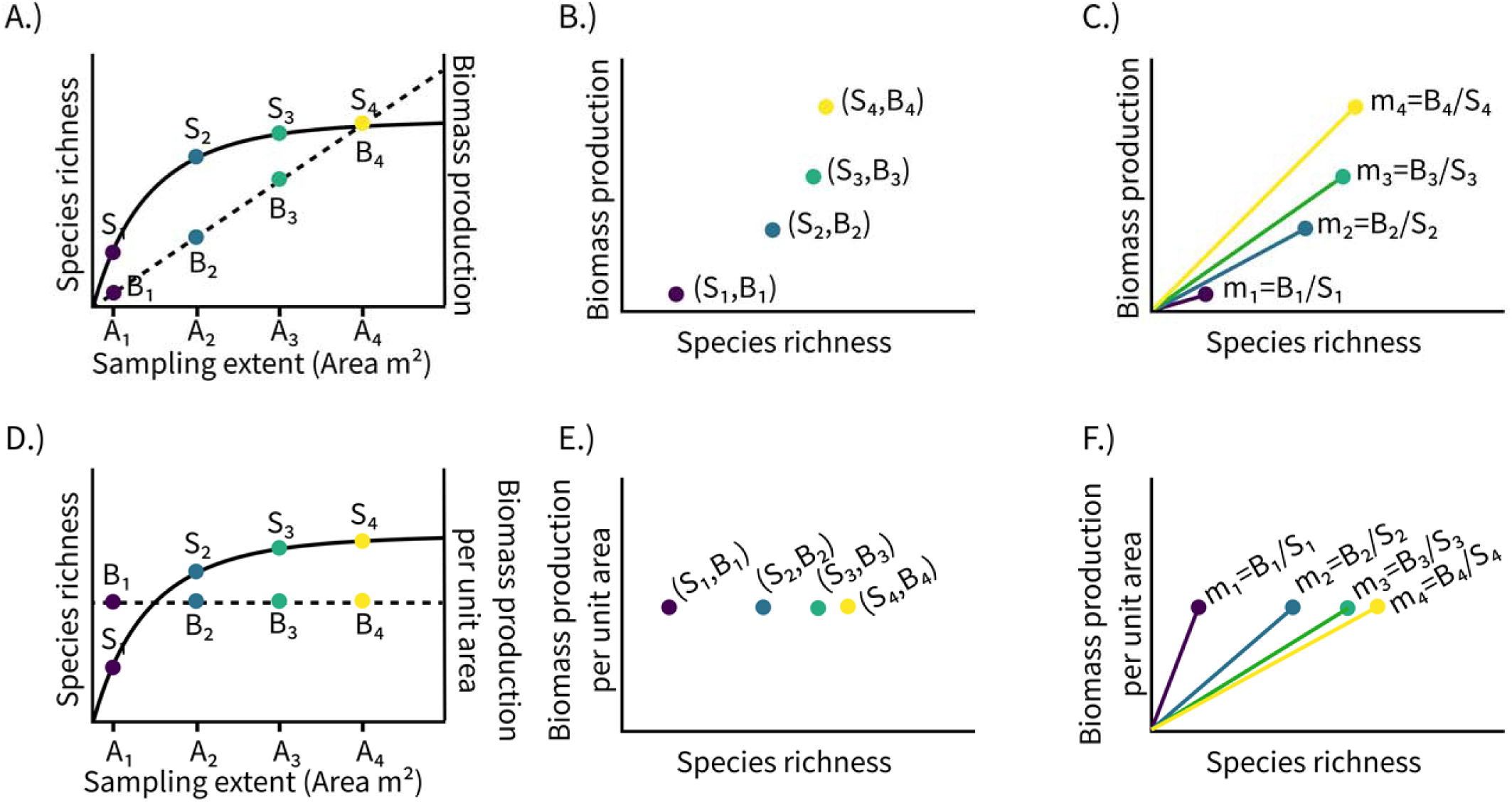
Theoretical mathematical constraints on the plant species richness-biomass relationship across different spatial scales. **A.)** Locally, species richness has a saturating relationship with area while biomass has a linear relationship with sampling extent. For a specific area A_i_, the species-area curve represents the average number of species S_i_ found on plots of area A_i_. Similarly, for a specific area A_i_ the biomass area curve yields the average total biomass B_i_ of species found on that area. The corresponding coordinate pairs (S_i_,B_i_) form the graph of the species richness biomass production relationship **(B)**. These coordinate pairs reveal an underlying accelerating nonlinear relationship between species richness and biomass production across scales. At any given scale the species richness biomass production curve contains the origin (because at zero species richness biomass production is zero and vice versa), as well as the point (S_i_,B_i_) of measured averages. If the species richness biomass relationship is linear (see Fig. S1 for alternative interpretations of this shape) and goes through these two points, then the predicted slope of the species richness-biomass relationship is the average biomass for that area divided by the average species richness for that area **(C.)** and the slope of the species richness biomass production relationship increases with increasing spatial scale of sampling. **D.)** When biomass is measured in terms of per unit area it is not likely to change with increasing sampling extent. **E.)** If this is the case, then the species-area curve determines the slope of the species richness-biomass relationship with increasing spatial scale **(F.).** Thus, we expect the per unit area version of the species richness-biomass relationship to have a decreasing slope with increasing area and that the slope will decrease at a decelerating rate.

## Species richness-biomass relationships match macroecological expectations

We produced species-area relationships (Figure 2) and biomass-area relationships (Figure 3) from (1) nested plots increasing in size in an herbaceous grassland at Cedar Creek Ecosystem Science Reserve and (2) using a resampling protocol on the tree community in the Barro Colorado Island (BCI) 50-hectare plot. Note that with regards to our results we use the term biomass to refer to our measures and biomass production to refer to the general ecosystem function. We distinguish between these two terms because while biomass is often a good proxy for biomass production, the latter requires a measure of turnover through time which we do not present here. We found that, as predicted species richness saturated with increasing sampling extent at both sites (Tables 1; Figure 2a,b). Biomass increased linearly with increasing sampling extent at both sites (Table 1; Figure 3a,c). By contrast, biomass per m^2^ was invariant to sampling extent at both sites (Table 1, Figure 3b,d).

**Figure 2.**
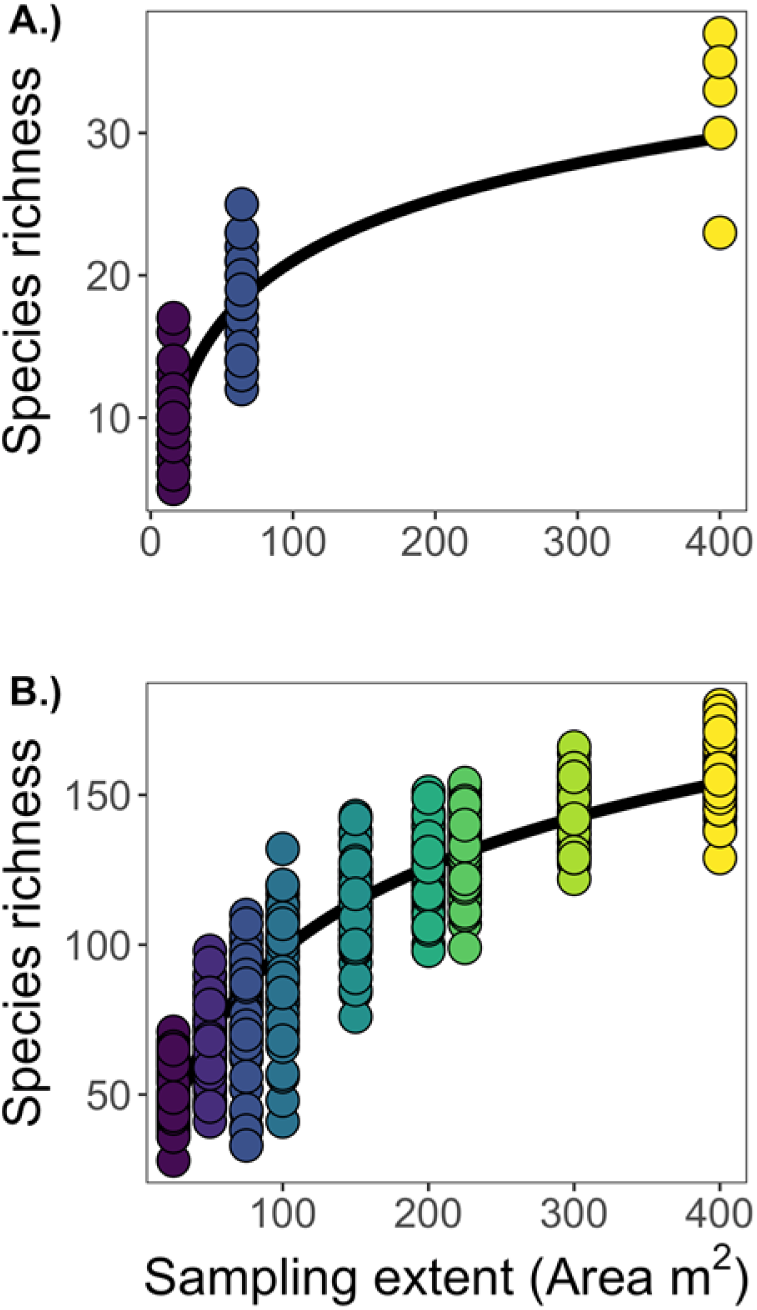
Species area curve in a Minnesota savanna and the Barro Colorado Island 50 Ha plot. **A.)** We found that plant communities in five recently burned savannas at the Cedar Creek Ecosystem Science Reserve had a saturating species-area curve as predicted. **B.)** Similarly, randomly sampled and combined sub-plots of the Barro Colorado Island 50 ha plot also had a significantly saturating species-area curve.

**Figure 3.**
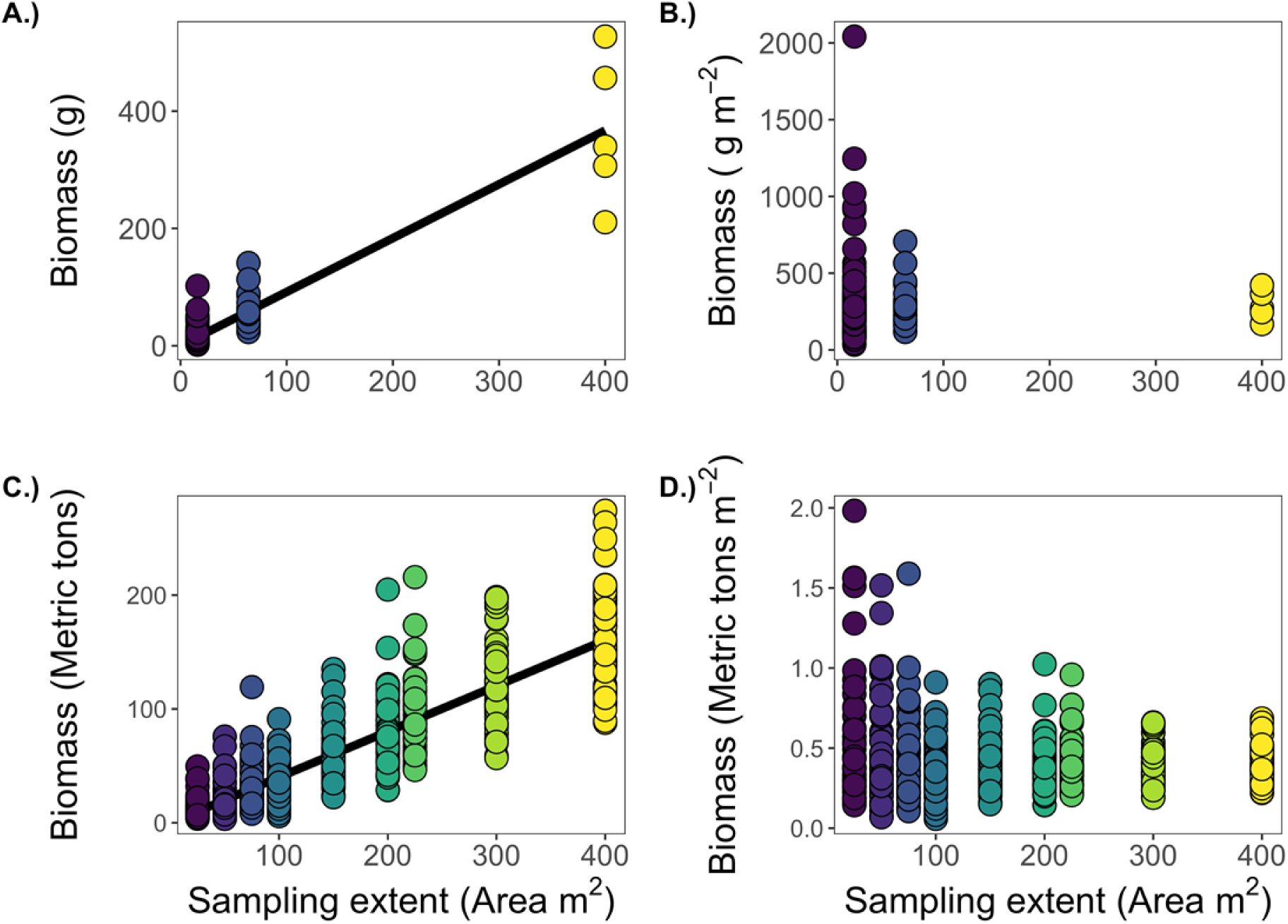
Biomass-area curves in a Minnesota savanna and the Barro Colorado Island 50-hectare plot. **A.)** When biomass was summed, it scaled linearly with increasing area. **B.)** When biomass was examined per m^2^ (as is commonly done in biodiversity-ecosystem function experiments), there was a linear trend that was not significant. **C.)** When biomass was summed in randomized subsamples of the BCI 50-ha plot, it increased in a linear fashion with increasing plot area. **D.)** Biomass per m^2^ in the BCI 50-ha plot did not significantly change with increasing plot area.

**Table 1.**
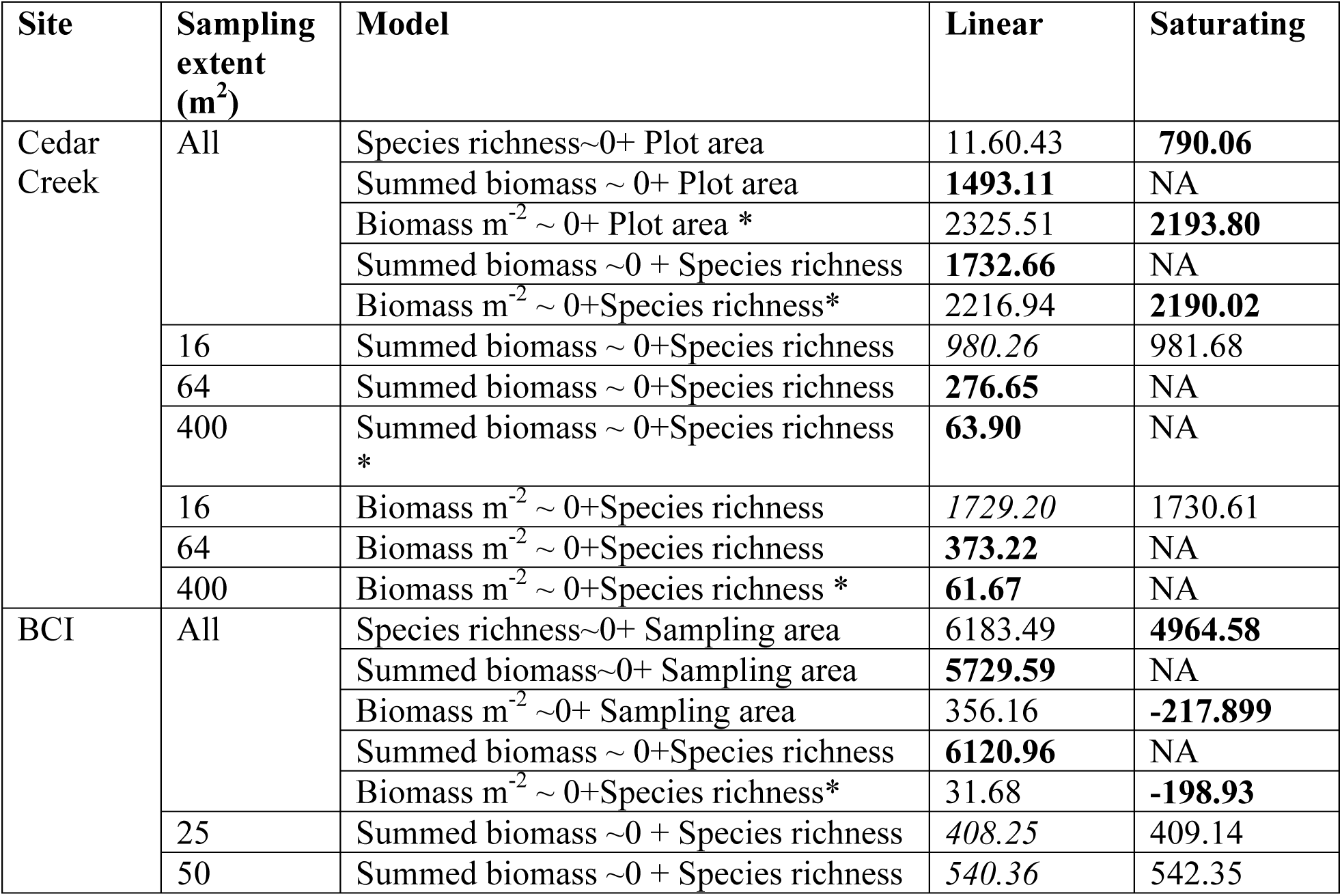
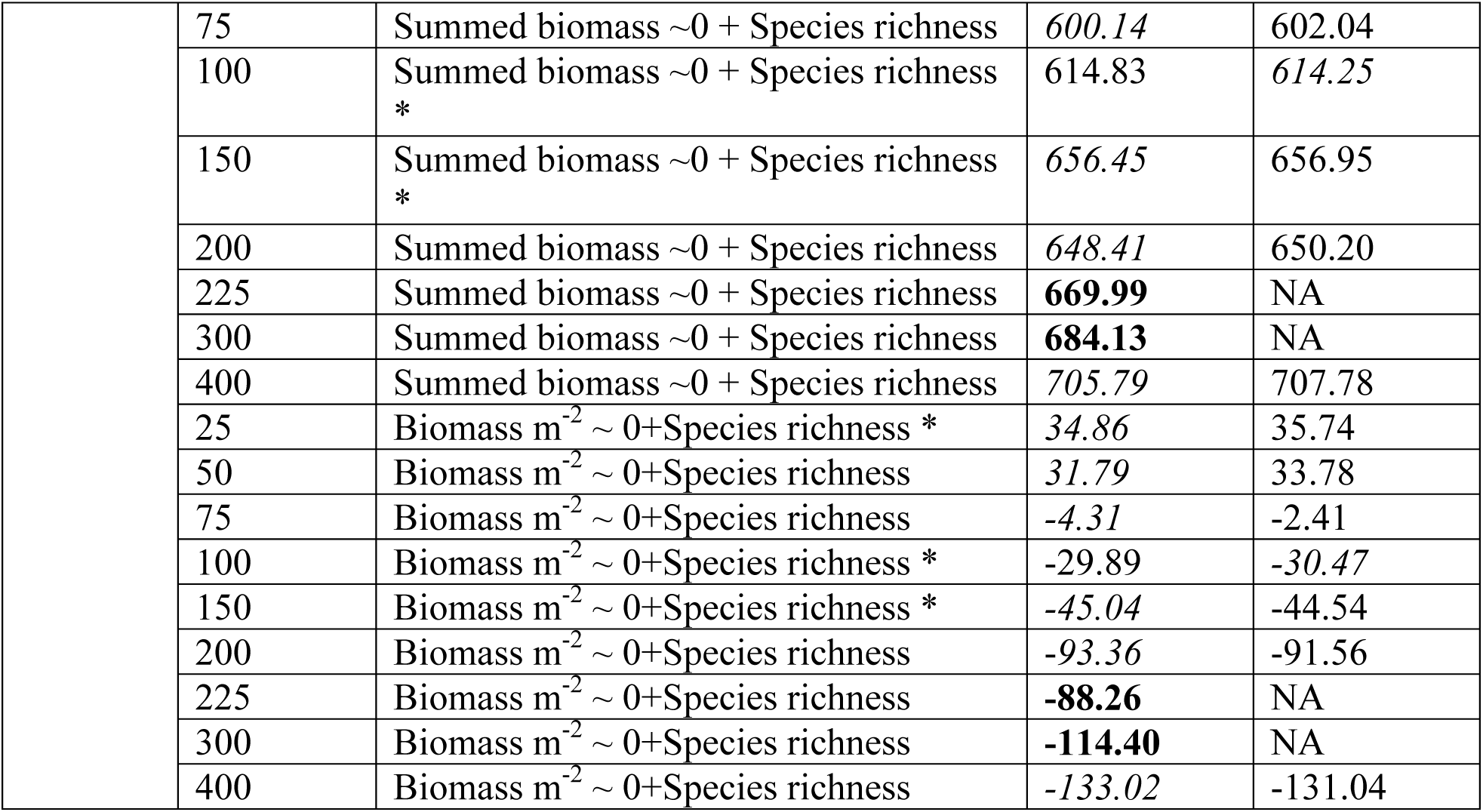
Akaike Information Criterion (AIC) for all implemented models at Cedar Creek Ecosystem Science Reserve (Cedar Creek) and Barro Colorado Island (BCI) . Numbers in bold are the best fit model (ΔAIC = 2). Numbers in italics are lower but not by a significant (>2) margin. For several models – a saturating model fit was unable to be fit using best practices for model parameterization resulting in no AIC value for saturating models for these individual models. * denotes a model where a simple linear model (dependent∼independent) was not significant indicating that there is no relationship between the independent and dependent variables. In this case, often a saturating model will be the best fit because of model artifacts rather than actual model fit when the intercept is included so we did not include the intercept in the linear model, denoted by ‘0+’ in the model.

We then used these area relationships to predict the slope of the species richness - biomass relationship at each sampling extent assuming that the species richness-biomass relationship intersects the origin (Figure 4). That is, we assumed that when an ecosystem has no species it also cannot produce biomass. At each sampling extent, we examined the species richness-biomass relationship in two different ways: (1) the relationship between total species richness per plot and total biomass per plot and (2) the relationship between total species richness per plot and biomass per m^2^(e.g., Isbell et al. 2011).

**Figure 4.**
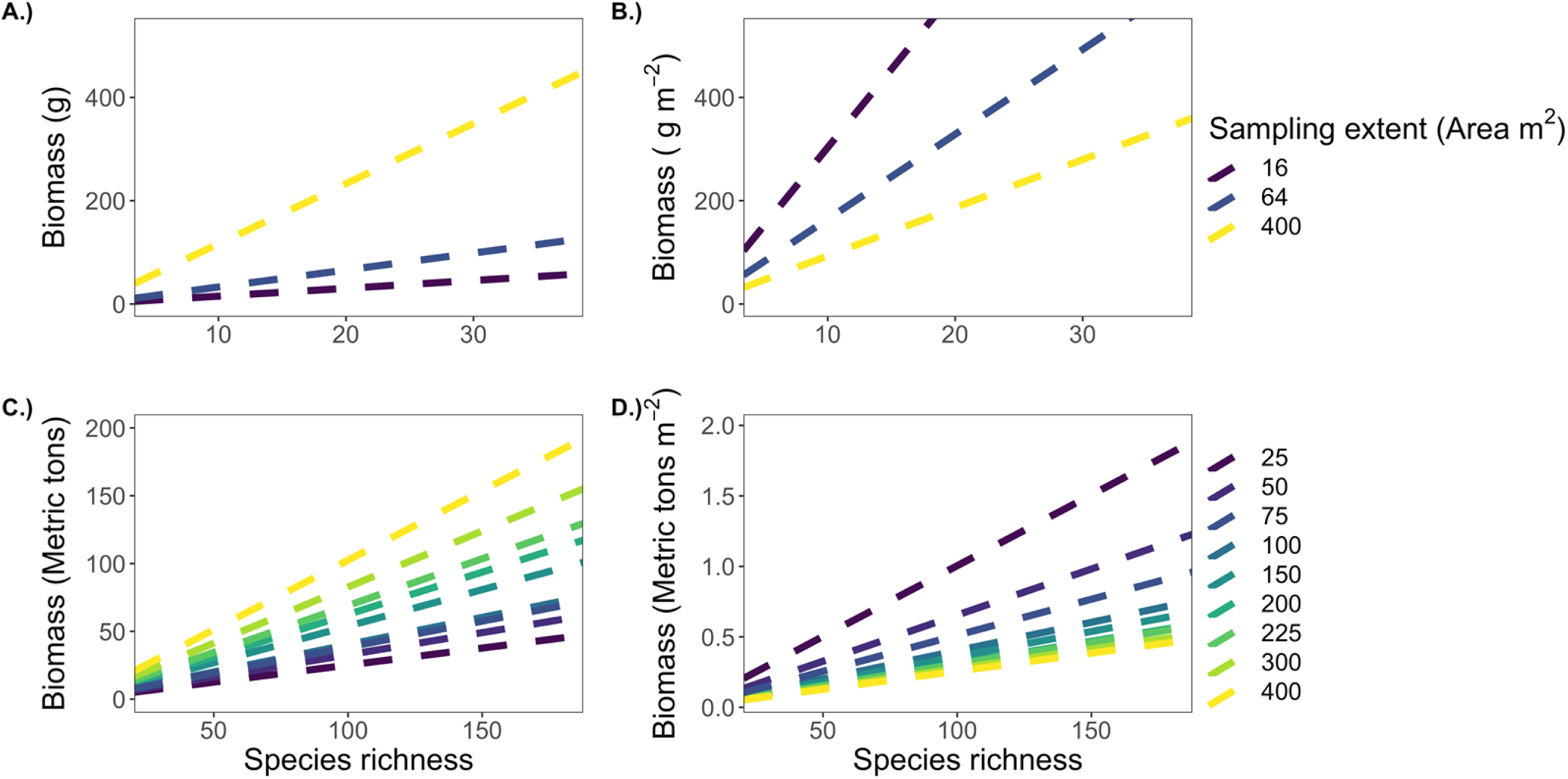
Predicted slopes of the species richness-biomass relationships at each scale for both absolute biomass and biomass per m^2^. When biomass is summed, we predict that the species richness-biomass production relationship will have an increasing slope with increasing area, for both recently burned savannas **(A.)** and the randomized subplots of the BCI 50-ha plot (**C.).** For biomass per m^2^, we predict that the slope of the species richness-biomass production relationship will decrease with increasing area for both recently burned savannas **(B.)** and the BCI 50-ha plot randomized subplots **(D.)**.

In accordance with our predictions, the slope of the species richness-total biomass relationship increased with increasing sampling extent at both sites (Figure 5a, c). In fact, no predicted slope for the species richness – biomass relationship was outside of the confidence limits of the observed slope (Table 2). Additional species contributed more to biomass production at the largest sampling extent, while the smallest sampling extent had the smallest relative gains in total biomass with each additional species. In terms of biomass per m^2^, we found that the slope of the species richness-biomass relationship decreased with increasing sampling extent (Figure 5b,d). That is, each additional species contributed less to biomass per m^2^ at the largest sampling extent. The smallest sampling extent had the largest relative gains in biomass per m^2^ with each additional species. At BCI, this decrease in slope was significant with increasing sampling extent, while at Cedar Creek it was not (Table 3).

**Figure 5.**
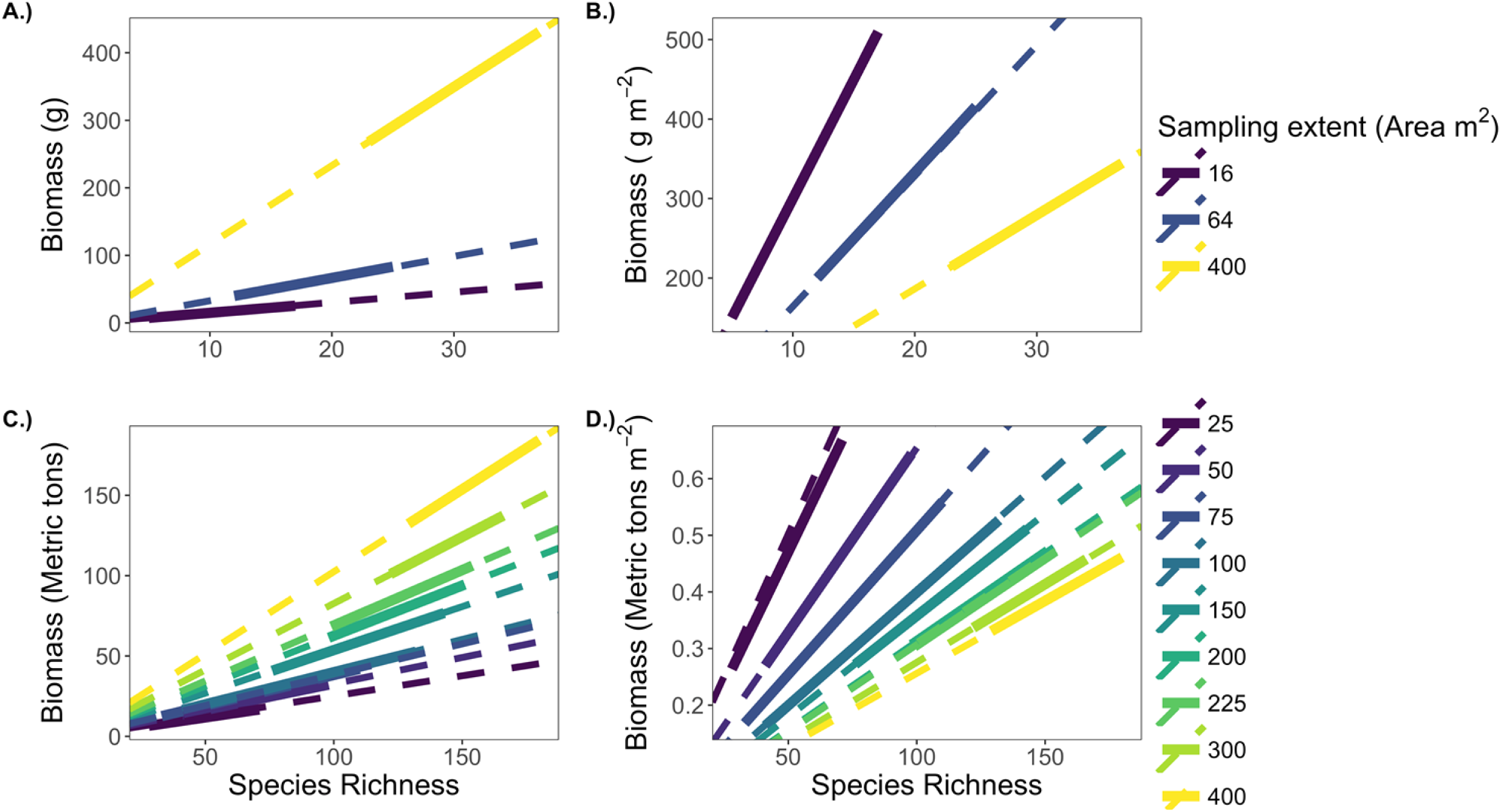
Actual species richness-biomass relationships and predicted species richness-biomass relationships in recently burned savannas and randomly selected subplots of the Barro Colorado Island 50-hectare plot. Throughout – dashed lines represent predictions, solid lines represent linear models from data. **A.)** When biomass is summed, the slope of the species richness-biomass production relationship increases with increasing spatial scale as predicted in five recently burned savannas. **B.)** When biomass was measured as biomass per m^2^, as is commonly calculated from biodiversity-ecosystem function experiments, the slope of the species richness-biomass production relationship decreases with increasing spatial scale. **C.)** Furthermore, from randomly subsampled subplots of the BCI 50-hectare plot, we found similarly that the species richness-biomass production (in terms of total aboveground biomass) relationship had an increasing slope with increasing spatial scale. **D.)** When biomass production was calculated as biomass per m^2^ from randomly subsampled subplots of the BCI 50-hectare plot, we found that the slope of the species richness-biomass production relationship decreased with increasing spatial scale.

**Table 2.** Predicted and actual slopes of species richness-biomass relationships. We used the species-area curve and biomass-area curves from a natural savanna at Cedar Creek Ecosystem Science Reserve and the forest inventory data from Barro Colorado Island to predict the slope of the species richness-biomass relationship at both sites across spatial scales. Our predictions were within the confidence limits of the observed slope at every spatial scale.

| Site | Biomass approach | Scale | Predicted Slope | Observed Slope | Lower Confidence Limit | Upper Confidence Limit |
| --- | --- | --- | --- | --- | --- | --- |
| BCI | Summed biomass | 25 | 0.25 | 0.24 | 0.19 | 0.3 |
|  |  | 50 | 0.32 | 0.33 | 0.28 | 0.38 |
|  |  | 75 | 0.35 | 0.34 | 0.3 | 0.39 |
|  |  | 100 | 0.43 | 0.42 | 0.36 | 0.47 |
|  |  | 150 | 0.55 | 0.56 | 0.5 | 0.61 |
|  |  | 200 | 0.65 | 0.65 | 0.59 | 0.71 |
|  |  | 225 | 0.66 | 0.66 | 0.6 | 0.71 |
|  |  | 300 | 0.82 | 0.82 | 0.76 | 0.87 |
|  |  | 400 | 1.01 | 1.01 | 0.96 | 1.07 |
| BCI | Biomass per square meter | 25 | 0.0102 | 0.0097 | 0.0077 | 0.0118 |
|  |  | 50 | 0.0064 | 0.0066 | 0.0056 | 0.0075 |
|  |  | 75 | 0.0047 | 0.0046 | 0.004 | 0.0052 |
|  |  | 100 | 0.0043 | 0.0042 | 0.0036 | 0.0047 |
|  |  | 150 | 0.0036 | 0.0037 | 0.0033 | 0.0041 |
|  |  | 200 | 0.0032 | 0.0032 | 0.003 | 0.0035 |
|  |  | 225 | 0.0029 | 0.0029 | 0.0027 | 0.0032 |
|  |  | 300 | 0.0027 | 0.0027 | 0.0025 | 0.0029 |
|  |  | 400 | 0.0025 | 0.0025 | 0.0024 | 0.0027 |
| Cedar Creek | Summed biomass | 16 | 1.51 | 1.5 | 1.28 | 1.71 |
|  |  | 64 | 3.2 | 3.32 | 2.84 | 3.81 |
|  |  | 400 | 11.65 | 11.65 | 7.44 | 15.84 |
| Cedar Creek | Biomass per square meter | 16 | 30.29 | 29.99 | 25.7 | 34.27 |
|  |  | 64 | 16.42 | 16.64 | 14.21 | 19.07 |
|  |  | 400 | 9.32 | 9.32 | 5.95 | 12.67 |

**Table 3.** Results of analysis of covariance tests to determine differences in the plant diversity-biomass production relationship across sampling extents at Cedar Creek Ecosystem Science Reserve (mixed effect model with nested random effects: (1|Site/Plot) as random factor) and at the Barro Colorado Island 50-ha plot. Results presented in bold are considered statistically significant (p<0.05).

| Site | Model parameters |  | Fixed effect parameters |  |  |
| --- | --- | --- | --- | --- | --- |
|  | Biomass approach | DF | Fixed effect | T | P value |
| Cedar Creek | Summed biomass | 148 | **Species richness** | **2.25** | **0.03** |
|  |  | 148 | Sampling extent | 0.84 | 0.40 |
|  |  | 148 | **Species richness\*sampling extent** | **2.41** | **0.017** |
|  | Biomass per square meter | 148 | Species richness | 1.54 | 0.125 |
|  |  | 148 | Sampling extent | -0.42 | 0.672 |
|  |  | 148 | Species richness*Sampling extent | 0.06 | 0.955 |
| BCI | Summed biomass | 617 | **Species richness** | **7.19** | **<0.001** |
|  |  |  | Sampling extent | 0.37 | 0.711 |
|  |  |  | **Species richness \* Sampling extent** | **7.79** | **<0.001** |
|  | Biomass per square meter | 617 | **Species richness** | **20.10** | **<0.001** |
|  |  |  | Sampling extent | -0.605 | 0.545 |
|  |  |  | **Species richness\*Sampling extent** | **-2.523** | **0.012** |

## Standardizing ecosystem functioning measures across sampling extents

We found that the slope of the species richness-total biomass relationship increased with increasing sampling extent and alternatively that the slope of the species richness-biomass per m^2^ relationship decreased with increasing sampling extent. The biomass per unit area (i.e. m^2^ or ha) approach, which is the typical method of comparison in biodiversity-ecosystem functioning studies (e.g., ^10, 12, 14, 17, 32, 33^), may not be particularly useful when comparing biodiversity-ecosystem functioning relationships across spatial scales. Reducing biomass to per unit area removes sampling extent from the biomass calculation and makes biomass a constant with respect to sampling extent while species richness continues to scale with increasing sampling extent. Further, because species richness scales non-linearly with increasing sampling extent, simply using a per unit area measure of species richness does not make biomass per m^2^ and species richness scale similarly.

## Applying macroecological predictions across scales, time, and to experimental approaches

A critical conclusion from our findings is that we can use macroecological patterns to accurately predict the scaling of the species richness-biomass production relationship at local spatial scales. However, the local to global scale species-area curve (both log transformed) is likely triphasic rather than saturating. As the ***log***(area) considered expands, the ***log***(species richness) saturates at local scales. At regional scales, the ***log*(** species richness) increases approximately linearly with increasing ***log*** (area). At landscape to global scales, the ***log***(species richness) and increases at an accelerating rate with increasing area ^26, 28, 30^. Given these underlying macroecological patterns, we expect that the scaling of the species richness-biomass relationship will also change as we move from local to global scales (when both species richness and biomass are log-transformed, Figure 6, see also^35^). In terms of ***log***(biomass production) (Fig. 6a), we expect that as ***log***(area) increases the slope of the relationship between the two will increase locally, stay the same regionally, and decrease slightly globally. Importantly, this global slope will not decrease below the slope of the highest local slope (Fig. 6b). That is, each additional species will contribute proportionally the most to biomass production at regional scales and the least at the smallest local scale. Alternatively, in terms of ***log***(biomass production per unit area), we expect that the species-area curve will change non-linearly from local to global scales and the ***log***(biomass production per unit area) will be invariant to scale (Fig. 6c). In this case, the slope of the relationship between species richness and biomass production will decline as area increases (Fig. 6d). That is, species will contribute the most to biomass production per unit area at local scales and the least globally.

**Figure 6.**
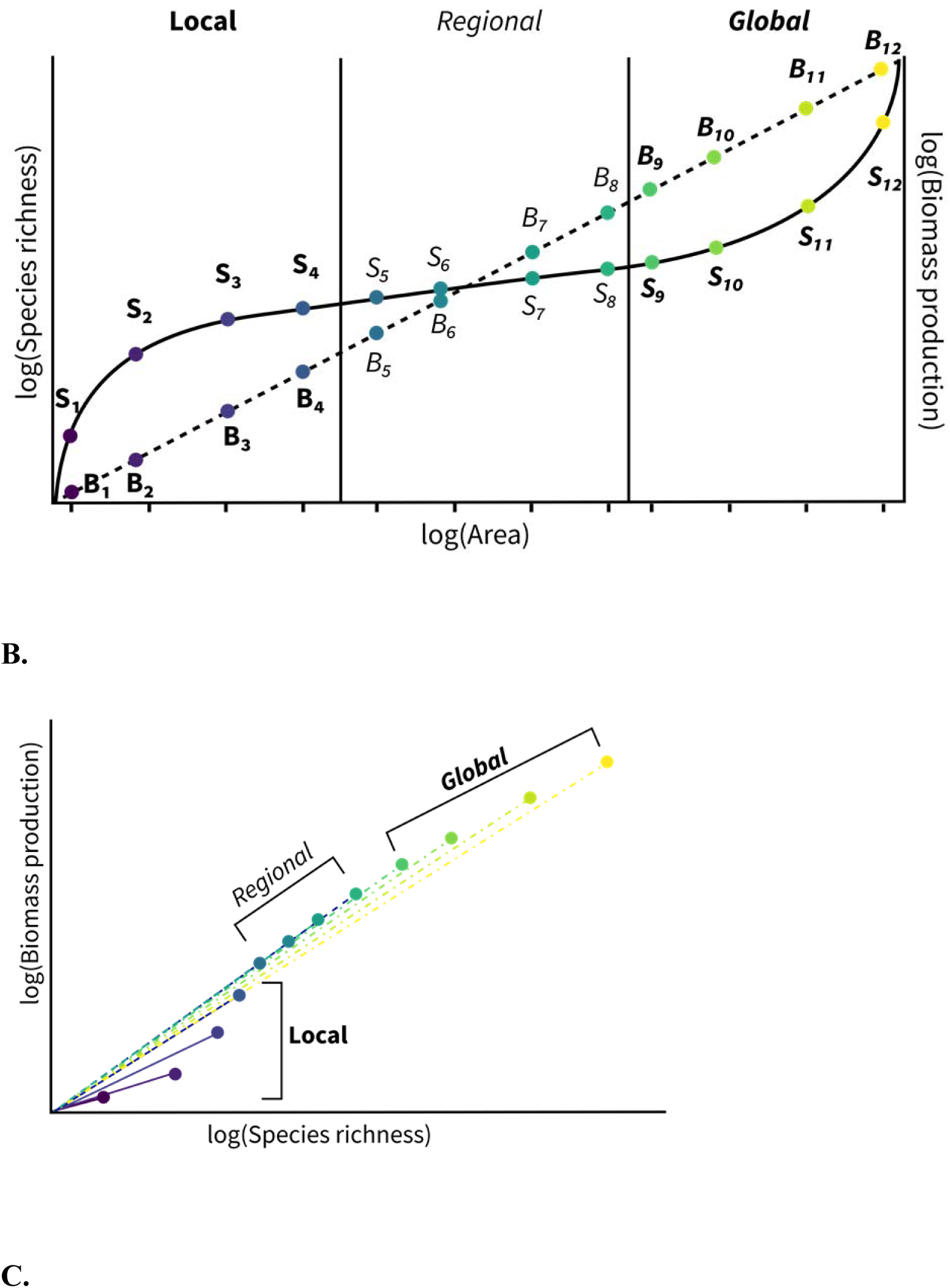
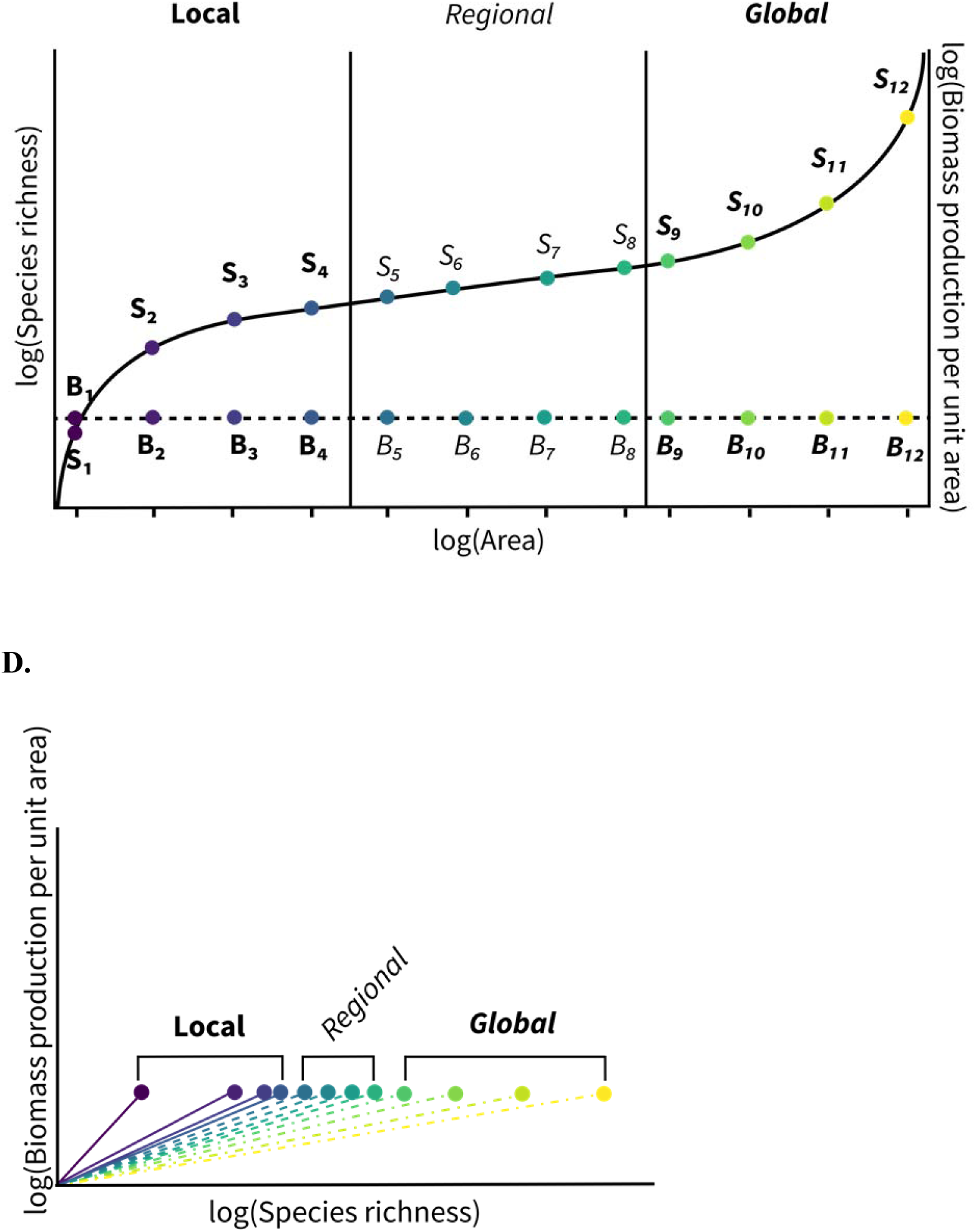
Graphical predictions for how the log(species richness)-log(biomass production) relationship will scale with increasing log(area) from local to regional and global scales depending on the way in which biomass production is calculated. A.) In log space, the species area curve is thought to be triphasic as log(area) increases from local to regional and then global scales (black curve). The log transformed total biomass production likely monotonically increases with increasing spatial scale (grey line). B.) The slope of the log(species richness)-log(biomass production) relationship increases with increasing spatial scale at local to the regional scales, does not change at regional scales, and decreases slightly at global scales. This would indicate that the exponents of the non-log transformed relationships increase with increasing spatial scale. C.) In log space, the species area curve is triphasic (black line). Log-transformed biomass per unit area (e.g. per m^2^ or per ha) is invariant to area. D.) The slope of the log(species richness)-log(biomass production) relationship decreases with increasing area at local, regional, and global scales. This would indicate that the exponents of the non-log transformed relationships decrease with increasing spatial scale

The general scaling relationship found here will also hold for any other analysis that combines a saturating relationship with a linear relationship. For example, the species richness-time relationship, while less well studied than the species-area relationship, also saturates at “local” time scales^26, 36, 37^. If we assume that biomass production per unit area remains relatively constant over time in an equilibrium community then we predict that the slope of the species richness-biomass relationship will decrease with increasing time and each additional species will be less important for biomass production per unit area with increasing time (Fig. S2a,b).

In cases where biomass production per unit area increases over time, our expectations would be the opposite. For example, in many biodiversity experiments, biomass per m^2^ increases through time (e.g.,^8, 14^, Fig 2c). Further, initial planted species richness is often reported in publications from biodiversity experiments (e.g.,^8, 14^). This initial species richness is constant over time. If species richness remains constant (or decreases) through time and biomass per unit area increases, then the slope of the relationship between the two will increase with increasing time, as found by Reich et al.^14^(Figure S2d). That is, in biodiversity experiments, we predict that over time each additional species will contribute more to biomass production while in natural systems that are not undergoing succession, we predict the opposite (Fig S2a,b). Entirely different mechanisms are at work in those two scenarios, so there is no theoretical conflict between the contrasting patterns. However, these contrasts would not emerge without the foundational mathematical underpinnings explored here.

## Conclusions

How biodiversity-ecosystem functioning relationships change with increasing spatial and temporal scale has critical implications for how biodiversity is managed to provide ecosystem services. Current conservation, restoration, and valuation efforts depend on our ability to scale from local experimental evidence to larger regional and even global scales^38, 39^. The current capacity for providing this upscaling is limited^40^. However, our results reveal the mathematical underpinnings of how biodiversity-ecosystem functioning relationships change with spatial scale. For any ecosystem function that scales linearly with increasing area, the contribution of each additional species will increase with increasing sampling extent due to the saturating species-area relationship. Further, all measures of biodiversity and all measures of ecosystem functioning have individual relationships with space and time. The shape of these underlying relationships determines how they scale. These underlying macroecological patterns, however, may be highly variable and driven by a variety of mechanisms. Understanding these underlying scaling relationships may allow us to make generalizable predictions for the consequences of species loss on any given biodiversity-ecosystem functioning relationship at global scales. Finally, understanding the mathematical underpinnings for these patterns allows us to focus more precisely on the mechanisms that may cause these relationships to diverge from our theoretical expectations.

## Methods

### Site Description: Cedar Creek Ecosystem Science Reserve

We conducted this study at Cedar Creek Ecosystem Science Reserve (CCESR) in central Minnesota in July of 2012. Our sites were located on Typic Udipsamment soils^41^ with a mean annual precipitation of 77.60± 4.57 cm (95% confidence interval for 1962-2012). CCESR consists of a matrix of prairies and savannas. We used two oak savannas (predominantly *Quercus macrocarpa*) of approximately equal type, age, and fire frequency. We chose oak savannas with similar burning and establishment histories that had little woody biomass and coarse woody debris to maximize similarity in nutrient availability and community composition between sites. The two oak savannas we chose were both burned in 2011 after the summer growing season, therefore all herbaceous biomass was considered to be the current year’s growth. Both savannas were burned frequently, 9 out of every 10 years and 4 out of every 5 years respectively, and were established 48 years ago from abandoned farmland ^42^. The dominant tree species at both sites were bur oak (*Quercus macrocarpa*), red oak (*Quercus rubra*), and black oak (*Quercus velutina*).

We established five 20 m by 20 m plots across these two oak savanna sites (three in one and two in the other). We avoided portions of the savannas with high amounts of woody biomass because many of the common woody savanna species are resistant to fire and we could not be certain that any woody biomass was indicative of this season’s growth. We established all plots > 3 m from any adult trees. Each plot was divided into 25 4 m x 4 m subplots.

### Biomass harvest: CCESR

To measure biomass, we harvested the aboveground vegetation in 10 cm x 50 cm strips in the center of each subplot in July of 2012. We harvested all vegetation strips within 6 days of each other to avoid seasonal and temporal variation in composition and biomass. We sorted the harvested vegetation to the species level when possible (95.46% of samples), genus level when species was not possible due to inadequate size of leaf matter (3.3% of samples), or categorized it as live litter when unidentifiable (1.24% of samples). We calculated species richness as the number of identified species in a clip strip. Vegetation was refrigerated when not being sorted. Once sorted, we placed each species in a paper envelope and dried all samples for at least 10 days in a 40°C drying oven. After 10-15 days, we weighed all samples and recorded the dry biomass.

### Site Description: Barro Colorado Island 50 Hectare Plot

We used data collected in the Barro Colorado Island 50-hectare plot from the 2015 census to examine whether total aboveground biomass in forests demonstrated similarly predictable patterns to aboveground biomass in grasslands (see ^43^ for data collection methods).

### Barro Colorado Island 50-hectare plot: Subsampling method

We assigned each 5 x 5 m subplot within the BCI 50 hectare plot a random number. We then randomly subsampled (using the command “sample”) these subplots to assemble non-spatially contiguous subsamples of the following areas: 25, 50, 75, 100, 150, 200 225, 300, and 400 m^2^. We randomly selected 70 of each subsample size for inclusion in our analysis. We used 70 as our sample size because it allowed for the inclusion of 400 m^2^ sampling areas which are comparable to the largest sampling area included in our data from Cedar Creek Ecosystem Science Reserve. We subsampled 70 of smaller spatial scales to prevent differences in sample size which may be relevant for our dataset from Cedar Creek Ecosystem Science Reserve where sample sizes varied from 125 at the smallest scale to 5 at the largest scale due to the nested nature of our sampling design. Further, we used spatially non-contiguous subsamples because the shape of the species-area curve changes between randomly sampled landscapes and landscapes where nested samples have been taken ^28^. Thus, to ensure that our results were not dependent upon the nested sampling schema used at CCESR, we subsampled randomly within the BCI 50-ha plot.

### Data analysis

To assess whether the relationship between species richness and biomass production was dependent on the spatial scale of measurement, we also examined the relationship of both species richness and biomass production individually with increasing plot area. We fit linear and “Michaelis-Menten” (saturating) models to our data and selected the model with the lower Akaike Information Criterion (AIC) to determine whether the relationships between biomass production and area, species richness and area, and species richness and biomass production were linear or saturating. When we were unable to calculate parameters for a Michaelis-Menten curve using the “SSmicmen” command in R and an “nls” command would not converge with uniform starting values (i.e. 1 for both parameters), we assigned an NA to the saturating model and considered the linear model to be a better fit.

We then used a mixed effects ANCOVA to determine whether the relationship between species richness and biomass production changed with increasing area of the plot using the command lme when random effects were necessary and gls when random effects were not necessary. We tested all models for heteroscedasticity using a Breusch-Pagan test^44^. When models had significant heteroscedasticity, we added an exponential variance structure to the model. All analyses were conducted in R Statistical Software and plotted using “ggplot2”.

## Acknowledgements

Funding for the collection of field data at Cedar Creek Ecosystem Science Reserve was provided by the National Science Foundation Grant DUE-1129056 to GP and IG for the University of Wisconsin-Milwaukee Undergraduate Research in Biology and Mathematics (UBM) program. Cedar Creek Ecosystem Science Reserve was funded by NSF Long Term Ecological Research funds DEB-8114302, DEB-811884, DEB-9411972, DEB-0080382, DEB-0620652, DEB-1234162. The BCI forest dynamics research project was founded by S.P. Hubbell and R.B. Foster and is now managed by R. Condit, S. Lao, and R. Perez under the Center for Tropical Forest Science and the Smithsonian Tropical Research in Panama. Numerous organizations have provided funding, principally the U.S. National Science Foundation, and hundreds of field workers have contributed. Funding for KB was provided by a German Centre for Integrative Biodiversity Research (iDiv) flexible pool grant for “Community Assembly and the Functioning of Ecosystems”. Funding for AW was also provided by CSULA startup funds. KB and AC are further supported by iDiv (German Research Foundation - FZT 118). This work was partially catalyzed by an NCEAS/LTER-NCO working group containing JC, AC, AM, LW, and AW funded by the NSF LTER program (DEB 1234162) and the LTER Network Communications Office (DEB-1545288). Group leaders are F. Isbell, J. Cowles, and L. Dee.

## Author contributions

KEB, IG, GP, SAS, and AJW designed the original version of this study. IG and GP provided funding for field work at Cedar Creek Ecosystem Science Reserve. JS, KY, KEB, and AJW planned and carried out field work. JS and KY produced initial drafts and analyses under the guidance of KEB, AJW, GP, and IG. KEB performed all subsequent analyses and rewrote the manuscript for submission under the guidance of AJW. All authors contributed substantially to the development of the concepts in the manuscript and revision of the text.

## Data availability statement

The data used from the BCI 50 ha plot are publicly available upon request at: ctfs.si.edu/webatlas/datasets/bci. Data collected at Cedar Creek Ecosystem Science Reserve will be made publicly available on Data Dryad prior to publication of this manuscript.

## Code availability statement

All code used for this analysis will be made publicly available with data on Data Dryad prior to the publication of this manuscript.

## Supplement

### ADDITIONAL RESULTS

#### Cedar Creek Ecosystem Science Reserve

We sampled 70 unique species in 125 vegetation strips in the two savannas. We found 12.55 g (± 2.09, 95% confidence intervals) of biomass and 7.74 (± 0.39) species on average in each sample. At the smallest sampling extent (4 m x 4 m, actual sampled area=0.05 m^2^), species richness ranged from 3 to 15 species. At the largest sampling extent (20 x 20 m, actual sampled area=1.25 m^2^), species richness ranged from 22 to 36 species. Indiangrass (*Sorghastrum nutans*) was the most abundant species representing 29.29% of all sampled biomass.

We found a significant and saturating species richness-area relationship and a significant and linear total biomass-area relationship (Table 1). There was no relationship between biomass m^-2^ and sampling extent (Table 1). The total biomass-species richness relationship was significant (t_1,148_ = 2.25, p=0.030, Table 3) and linear at all sampling extent (though not significantly for 16 m^2^ plots). Further, the sampling extent significantly increased the slope of the summed biomass-species richness relationship as predicted (t_1,148_ = 2.41, p=0.017, Table 3). Increasing species richness did not increase biomass m^-2^ (t_1,148_ = 1.54, p=0.125, Table 3) and the sampling extent did not significantly alter the biomass m^-2^-species richness relationship (t_1,148_ = 0.06, p=0.955, Table 3).

#### Barro Colorado Island 50 ha plot

Similar to CCESR, we found a saturating relationship between species richness and the sampling extent as predicted and the summed biomass increased linearly with increasing sampling extent (Table 1). The species richness-summed biomass relationship was significant (t_1,617_= 7.17, p<0.001, Table 3) and was linear at all sampling extents except for 100 m^2^ (Table 1). However, the linear model was only a significantly better fit than the saturating one at the 225 m^2^ and 300 m^2^ sampling extents where the saturating model did not converge (Table 1). Further, the sampling area significantly increased the slope of the species richness-biomass relationship as predicted (t_1,617_ = 7.79, p<0.001, Table 3). Similarly, the biomass per m^2^-species richness relationship was significant (t_1,617_= 20.10, p<0.001, Table 3) and linear at all sampling extents except 100 m^2^. Furthermore, the linear model was only significantly better than the saturating model at the 225 m^2^ and 300 m^2^ sampling areas where the saturating model did not converge. With increasing sampling extent, the slope of the species-richness biomass relationship significantly decreased (t_1,617_ = -2.523, p=0.012, Table 3).

Further, at every sampling extent, at both sites the predicted slope based on the species-area curve and the biomass-area curve was within the confidence limits of the observed slope of the species richness-biomass relationship (Table 2).

**Figure S1:**
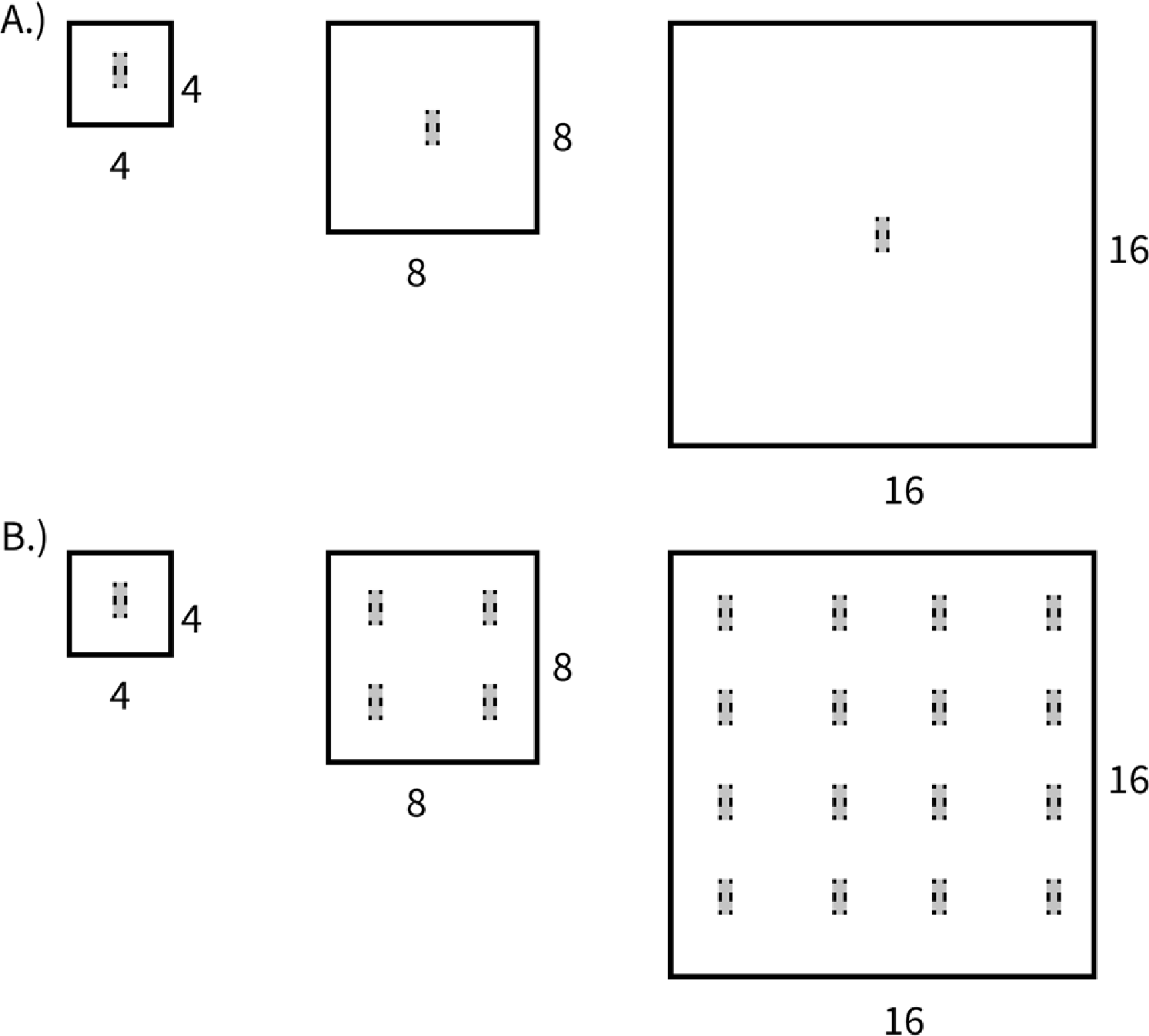
Sampling schematics in plots of three different spatial scales. Several studies have examined whether the species richness-biomass production (or the biomass production-species richness) relationship changes with increasing spatial scale. These papers use two sampling approaches. A.) Studies that measure the effect of spatial scale of the area *per se* on a sample of the same size take the same number of samples (grey boxes) from contiguous areas that are increasingly large. This type of study measures how the size of the area affects a sample which does not vary in size and may be helpful in studying edge effects or the pure effect of plot size in experimental settings (see Roscher et al. 2005). B.) Studies where the actual sampled area and thus sampling extent increases (this study). Here, sampling extent can increase in two ways. First, sampling can be non-destructive in which case an entire area is often sampled. Second (as pictured), if sampling is destructive, randomly spaced or evenly spaced subsamples may be taken. If this is the case, our examples require that this be proportional to the amount of area increased. For example, 1 sample is taken in a 4 x 4 m plot. In an 8 x 8 m plot, the area increases by a factor of 4. Therefore, in the 8 x 8 m plot we take 4 samples.

**Figure S2.**
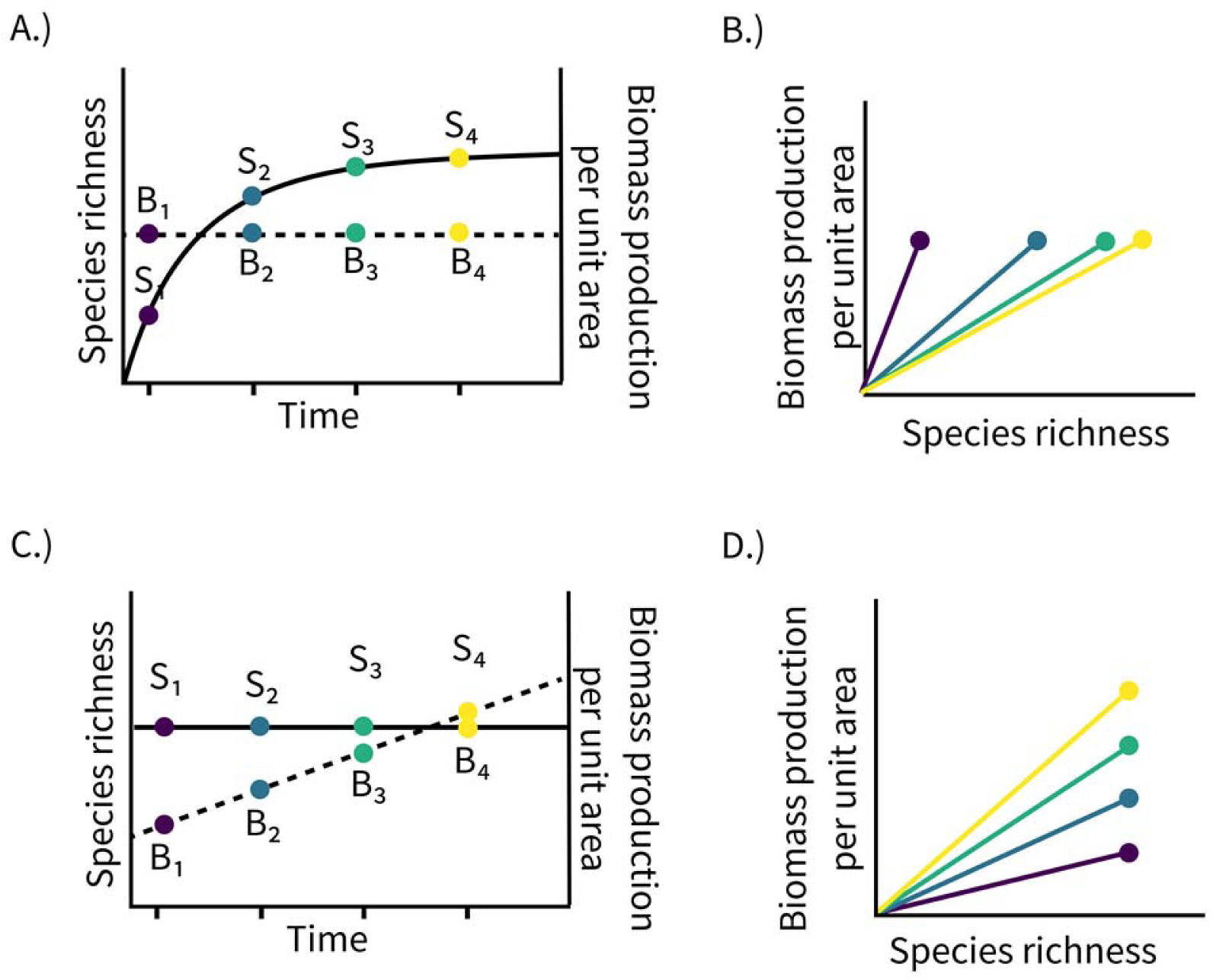
Graphical predictions for how species richness-biomass production relationships scale with time in natural vs. experimental systems. A.) In natural systems, the sampled species richness increases with increasing study time until saturation while the biomass production per unit area remains relatively constant. B.) Combining a saturating species-time relationship with a constant biomass production per unit area means that each additional species will contribute proportionally less with increasing time. C.) In biodiversity experiments, average species richness is held constant over time and biomass production per unit area increases after the establishment of the experiment. D.) Combining a constant species-time relationship with a linearly increasing biomass production per unit area means that each individual species will matter more with increasing time from the establishment of the experiment.

## References

1. Millenium Ecosystem Assessment Ecosystem and human well-being: synthesis. (Island Press, 2005).

2. Newbold, T. et al. Global effects of land use on local terrestrial biodiversity. Nature 520, 45–50 (2015).

3. Tittensor, D. P. et al. A mid-term analysis of progress toward international biodiversity targets. Science 346, 241–244 (2014).

4. Pereira, H. M. et al. Scenarios for Global Biodiversity in the 21st Century. Science 330, 1496–1501 (2010).

5. Pimm, S. L. et al. The biodiversity of species and their rates of extinction, distribution, and protection. Science 344, 1246752 (2014).

6. Hooper, D. U. et al. Effects of Biodiversity on Ecosystem Functioning: A Consensus of Current Knowledge. Ecol. Monogr. 75, 3–35 (2005).

7. Cardinale, B. J. et al. Biodiversity loss and its impact on humanity. Nature 486, 59– 67 (2012).

8. Weisser, W. W. et al. Biodiversity effects on ecosystem functioning in a 15-year grassland experiment: Patterns, mechanisms, and open questions. Basic Appl. Ecol. 23, 1–73 (2017).

9. Tilman, D. et al. Diversity and Productivity in a Long-Term Grassland Experiment. Science 294, 843–845 (2001).

10. Reich, P. B. et al. Plant diversity enhances ecosystem responses to elevated CO2 and nitrogen deposition. Nature 410, 809–810 (2001).

11. Van Ruijven, J. & Berendse, F. Positive effects of plant species diversity on productivity in the absence of legumes. Ecol. Lett. 6, 170–175 (2003).

12. Roscher, C. et al. The role of biodiversity for element cycling and trophic interactions: an experimental approach in a grassland community. Basic Appl. Ecol. 5, 107–121 (2004).

13. Schnitzer, S. A. et al. Soil microbes drive the classic plant diversity-productivity pattern. Ecology 92, 296–303 (2011).

14. Reich, P. B. et al. Impacts of biodiversity loss escalate through time as redundancy fades. Science 336, 589–592 (2012).

15. Zhang, Y., Chen, H. Y. & Reich, P. B. Forest productivity increases with evenness, species richness and trait variation: a global meta-analysis. J. Ecol. 100, 742–749 (2012).

16. Isbell, F. et al. Biodiversity increases the resistance of ecosystem productivity to climate extremes. Nature 526, 574–577 (2015).

17. Duffy, J. E., Godwin, C. M. & Cardinale, B. J. Biodiversity effects in the wild are common and as strong as key drivers of productivity. Nature 549, 261–264 (2017).

18. van der Plas, F. Biodiversity and ecosystem functioning in naturally assembled communities. Biol. Rev. 0, (2019).

19. Cardinale, B. J. et al. The functional role of producer diversity in ecosystems. Am. J. Bot. 98, 572–592 (2011).

20. Hooper, D. U. et al. A global synthesis reveals biodiversity loss as a major driver of ecosystem change. Nature 486, 105–108 (2012).

21. McGill, B. J., Dornelas, M., Gotelli, N. J. & Magurran, A. E. Fifteen forms of biodiversity trend in the Anthropocene. Trends Ecol. Evol. 30, 104–113 (2015).

22. Gleason, H. A. On the relation between species and area. Ecology 3, 158–162 (1922).

23. Braun-Blanquet, J. Plant Sociology. The study of plant communities. Plant Sociol. Study Plant Communities First Ed (1932).

24. Cain, S. A. The species-area curve. Am. Midl. Nat. 573–581 (1938).

25. Rice, E. L. & Kelting, R. W. The Species--Area Curve. Ecology 36, 7–11 (1955).

26. Preston, F. W. Time and Space and the Variation of Species. Ecology 41, 611–627 (1960).

27. Connor, E. F. & McCoy, E. D. The statistics and biology of the species-area relationship. Am. Nat. 791–833 (1979).

28. Rosenzweig, M. L. Species Diversity in Space and Time. (Cambridge University Press, 1995).

29. Lomolino, M. V. Ecology’s most general, yet protean 1 pattern: the species-area relationship. J. Biogeogr. 27, 17–26 (2000).

30. Lomolino, M. v. The species–area relationship: new challenges for an old pattern. Prog. Phys. Geogr. 25, 1–21 (2001).

31. Detto, M. & Muller-Landau, H. C. Fitting Ecological Process Models to Spatial Patterns Using Scalewise Variances and Moment Equations. Am. Nat. 181, E68–E82 (2013).

32. Spehn, E. M. et al. Ecosystem Effects of Biodiversity Manipulations in European Grasslands. Ecol. Monogr. 75, 37–63 (2005).

33. Isbell, F. et al. High plant diversity is needed to maintain ecosystem services. Nature 477, 199–202 (2011).

34. Meyer, S. T. et al. Effects of biodiversity strengthen over time as ecosystem functioning declines at low and increases at high biodiversity. Ecosphere 7, e01619 (2016).

35. Thompson, P. L., Isbell, F., Loreau, M., O’Connor, M. I. & Gonzalez, A. The strength of the biodiversity–ecosystem function relationship depends on spatial scale. Proc R Soc B 285, 20180038 (2018).

36. Adler, P. B. & Lauenroth, W. K. The power of time: spatiotemporal scaling of species diversity. Ecol. Lett. 6, 749–756 (2003).

37. White, E. P. et al. A comparison of the species–time relationship across ecosystems and taxonomic groups. Oikos 112, 185–195 (2006).

38. Isbell, F. et al. Linking the influence and dependence of people on biodiversity across scales. Nature 546, 65 (2017).

39. Naidoo, R. et al. Global mapping of ecosystem services and conservation priorities. Proc. Natl. Acad. Sci. 105, 9495–9500 (2008).

40. Bockstael, N. E., Freeman, A. M., Kopp, R. J., Portney, P. R. & Smith, V. K. On Measuring Economic Values for Nature. Environ. Sci. Technol. 34, 1384–1389 (2000).

41. Dickie, I. A., Schnitzer, S. A., Reich, P. B. & Hobbie, S. E. Is oak establishment in old-fields and savanna openings context dependent? J. Ecol. 95, 309–320 (2007).

42. Peterson, D. W. & Reich, P. B. Prescribed Fire in Oak Savanna: Fire Frequency Effects on Stand Structure and Dynamics. Ecol. Appl. 11, 914–927 (2001).

43. Condit, R. Tropical forest census plots: methods and results from Barro Colorado Island, Panama and a comparison with other plots. (Springer Verlag, 1998).

44. Breusch, T. S. & Pagan, A. R. A Simple Test for Heteroscedasticity and Random Coefficient Variation. Econometrica 47, 1287–1294 (1979).

